# Astrocytes tune neuronal excitability through the Ca^2+^-activated K^+^ current sIAHP

**DOI:** 10.1101/2024.01.24.576973

**Authors:** Sara Expósito, Samuel Alberquilla, Eduardo D. Martín

## Abstract

Neurons have the unique ability to integrate synaptic information by modulating the function of the voltage-gated membrane ion channels, which govern their excitability. Astrocytes play active roles in synaptic function, from synapse formation and maturation to plasticity processes. However, it remains elusive whether astrocytes can impact the neuronal activity by regulating membrane ion conductances that control the intrinsic firing properties. Here, we found that astrocytes, by releasing adenosine, enhance the slow Ca^2+-^activated K^+^ current (sIAHP) in CA1 hippocampal pyramidal neurons. Remarkably, we showed that interneuron activity was involved in the astrocyte-mediated sIAHP modulation. Indeed, both synaptically activated and optogenetically stimulated hippocampal interneurons evoked coordinated signaling in astrocytes and pyramidal neurons, which relied on GABA_B_ and adenosine A1 receptors activation. In addition, the selective genetic ablation of GABA_B_ receptors in CA1 astrocytes prevented the spike frequency adaptation in pyramidal cells after interneuron activation. Therefore, our data reveal the astrocyte capability to modulate the intrinsic membrane properties that dictate neuronal firing rate and hippocampal networks activity.

## INTRODUCTION

The functional response of neurons results from the integration of synaptic inputs and intrinsic membrane properties. There is large variation in the intrinsic properties among neurons in different brain regions, or even into the same area. In the hippocampus, action potentials in pyramidal neurons are followed by a prolonged after hyperpolarization (AHP) through a Ca^2+^-dependent K^+^-current (IAHP), which is largely activated following Ca^2+^ influx via voltage-gated Ca^2+^ channels^1^. IAHP has a critical role in neuronal excitability, controlling action potential shaping and firing patterns^2–4^. Three AHP components are widely described in rodent and humans^2,5–8^. The fast component is Ca^2+^ and voltage-dependent current and is activated immediately after action potentials^9,10^. The medium and slow components are voltage-independent but rely on cytosolic Ca^2+^ levels^9,10^. AHP components are mediated by Ca^2+^-activated K^+^ currents with fast (fIAHP), medium (mIAHP) and slow (sIAHP) decay kinetics^11^. The mIAHP and sIAHP provide a negative feedback control of neuronal excitability and spike frequency adaptation^12^, and it has been recognized that both components contribute to epileptogenesis^13–15^. In addition, sIAHP can be modulated by several neurotransmitters such as norepinephrine, serotonin, histamine, acetylcholine, glutamate, dopamine^15–18^ and hormones, such as estradiol^19^, revealing the complex regulatory mechanisms controlling this conductance.

Astrocytes are directly involved in synaptic physiology^20^ being a key player in tripartite synapse^21^. They respond to different neurotransmitters through functional receptors^22–32^ and they can release different gliotransmitters, through Ca^2+^-dependent and Ca^2+^-independent mechanisms^33–36^, such as glutamate, GABA (γ-aminobutyric acid) and ATP/adenosine, which activate neuronal receptors^32,37–40^. Furthermore, astrocytes are able to decode neuronal activity and adjust their responses to modulate synaptic transmission, by releasing glutamate or ATP/adenosine^41^, leading to a potent excitatory or inhibitory neuromodulation. In addition to the astrocytic ability to modulate synaptic transmission and plasticity, limited experimental evidence indicates that these glial cells regulate neuronal excitability. Astrocytes increase interneuron activity by closing K2P channels through the activation of P2Y1 receptors and decrease pyramidal neuron activity by opening GIRK channels through the activation of adenosine A_1_ receptors (A_1_R) in hippocampus^42^, and myelinated axons^43^. However, it remains unknown whether astrocytes can impact the sAHP potassium currents.

In the present study, we found that selective chemogenetic activation of hippocampal astrocytes by hM3Dq DREADD (Designer Receptors Exclusively Activated by Designer Drugs), which stimulates intracellular Ca^2+^ signaling and gliotransmission^44^, decreases neuronal excitability by increasing the sIAHP in CA1 pyramidal neurons. This enhancement of sIAHP involved the activation of G_i/o_ proteins/PLC-dependent pathway^41,45–47^. Both pharmacological and genetic approaches revealed that GABAergic interneurons trigger astrocytic Ca^2+^ elevations that stimulate the release of ATP/adenosine from astrocytes and the activation of A_1_R in pyramidal neurons. Therefore, these novelty data show how astrocytes control the intrinsic CA1 pyramidal neuron excitability by acting on sIAHP, regulating the firing properties of pyramidal neurons according to the hippocampal network demand/activity, and highlight the critical functional role of astrocytes as part of a multicellular complex signaling that governs hippocampal excitability.

## RESULTS

### Astrocytes decrease CA1 pyramidal neurons excitability by increasing sIAHP

In order to evaluate the potential role of astrocytic gliotransmission on CA1 intrinsic excitability, we used a DREADD approach to activate intracellular Ca^2+^ signaling in astrocytes. Mice were injected with AAV5-GFAP-hM3D(Gq)-mCherry and the genetically encoded Ca^2+^ indicator AAV5-gfaABC1D-cyto-GCaMP6f to activate and monitor Ca^2+^ levels in hippocampal astrocytes (Figure 1A, top). Ex vivo hippocampal recordings showed that local application of CNO (see Methods), a specific agonist of hM3Dq, transiently increased the Ca^2+^ event frequency of Gq-DREADD expressing astrocytes (Figure 1A, bottom; B; C), confirming its ability to selectively engage astrocytic signaling. Then, current-clamp (CC) whole-cell recordings were performed in CA1 pyramidal neurons in the area of transfected astrocytes (Figure 1D, top). To ensure reliable astrocytic network activation between different experiments, CNO (10 µM) was perfused through the slice. In these conditions, selective stimulation of Gq-DREADD expressing astrocytes significantly decreased the action potential firing rate (Figure 1D, bottom) and increased the spike-frequency adaptation (Figure 1E), measured as the progressive slowdown of the frequency of discharge after the first burst of spikes of a train.

**Figure 1:**
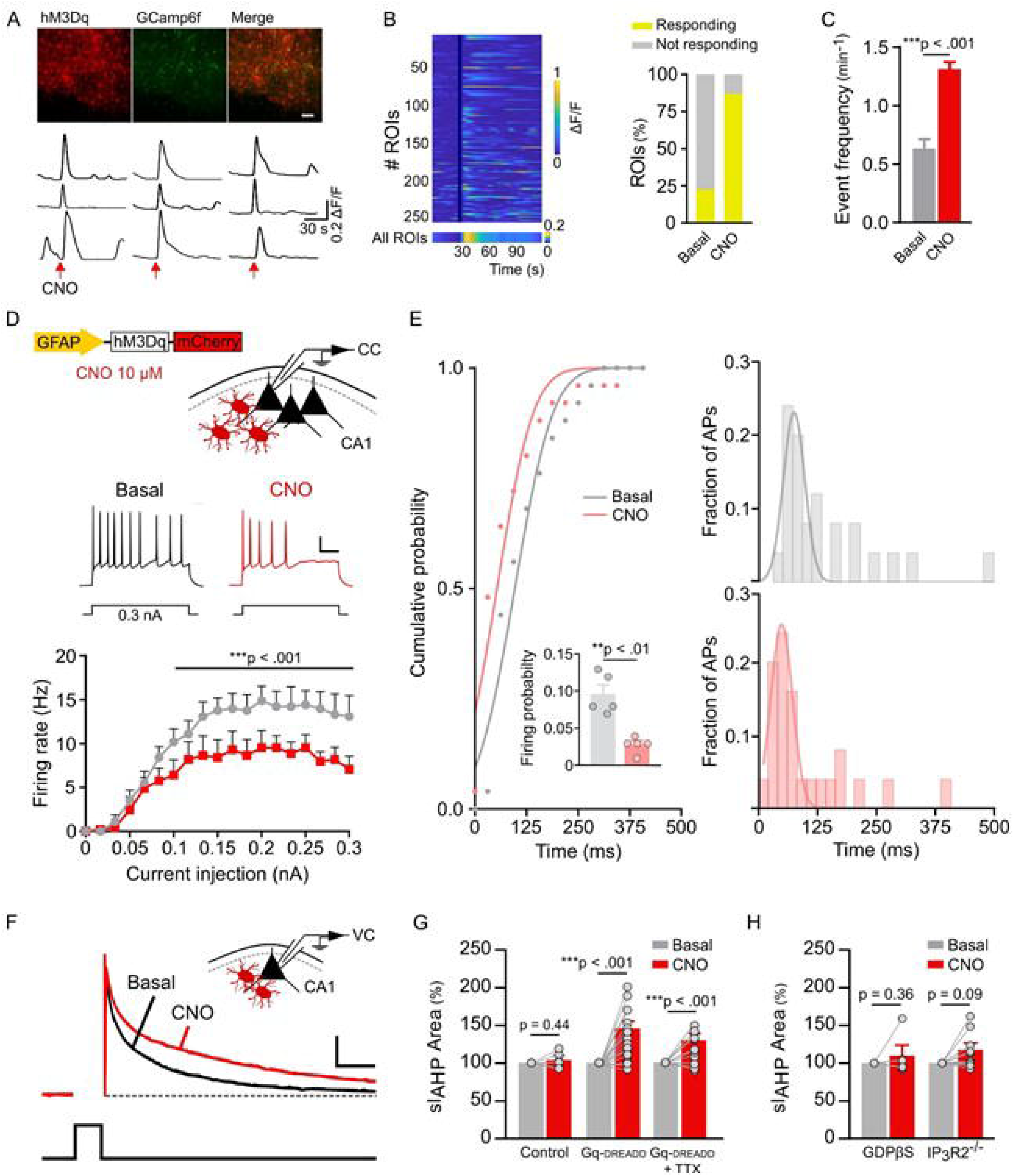
Astrocytes activation modulate CA1 pyramidal neurons firing frequency and spike accommodation by upregulation of sIAHP. **A**, Top: AAV5-GFAP-hM3Dq-mCherry and AAV5-gfaABC1D-cyto-GCaMP6f expression in astrocytes in the CA1 region of the hippocampus. Scale bar: 50 μm. Bottom: Representative Ca^2+^ traces showing an increase in astrocytic Ca^2+^ signaling after application of CNO (1 mM) (astrocytes stimulation denoted by red arrow). **B**, Left: Raster plot of ROIs activity, color coded according to fluorescence change (n=253). Right: Percentage of astrocytes showing any event (Responding) prior to the local application of CNO (basal, n=53) and first 60 seconds after applying the drug (CNO, n=146). **C**, Mean responses for the frequency of somatic Ca^2+^ fluctuation events (15 slices from three mice; Friedman test analysis, post hoc comparison with Dunn’s test). **D**, Top: Viral construct used and schematic drawing of the experimental design. Bottom: Representative traces of current-clamp (CC) recordings and summary data show a significant decline of pyramidal neuron firing frequency after astrocyte stimulation (10 µm CNO) in response to somatic current depolarizing pulses of 500 ms (n=9; two-way ANOVA, F(1, 320) = 49,69; post hoc comparison with Bonferroni test). Scale bar: 20 mV, 0.1 s. **E**, Left: Cumulative probability plot depicting the distribution of action potentials along the 500 ms of the depolarizing pulse. Inset: To study whether astrocyte activation prompts spike frequency adaptation, the probability that a neuron fires an action potential during the last 100 ms of the pulse was also analyzed. Here we show that, as adaptation is so marked after CNO perfusion, only a small number of currents are initiated so the firing probability declines significantly (n=5; Paired *t-*test). Right: The plot reveals that astrocyte activation results in a significant increase of spike frequency accommodation. As shown, neurons respond with an initial burst of spikes following CNO perfusion, but then remain more silent for the duration of the pulse than in their basal condition (25 bins of 20 ms each; Kolmogorov-Smirnov test, P=0.006). **F,** Superimposed voltage-clamp (VC) recordings of the sIAHP before and after CNO perfusion (10 μm) evoked by 200 milliseconds depolarizing pulses to 20 mV (bottom trace). Scale bar: 100 pA, 0.25 s. **G,** Astrocyte activation evokes sIAHP enhancement in Gq-DREADD animals (n=6 neurons for Control; n=18 for Gq-DREADD), also in presence of TTX (n=15). Paired *t-*tests. **H**, No effect upon sIAHP area was observed after perfusion of CNO over astrocytes with GDPβS-ablated signaling (n=6; Paired *t-*test) and IP3R2^-/-^ mice (n=11; Mann-Whitney test). Data are represented as mean ± SEM.

It is well established that sIAHP provides a strong negative feedback control of neuronal excitability^12^ and is responsible for the late phase of spike-frequency adaptation^48^. Therefore, the next step was to perform voltage-clamp (VC) recordings to evoke the sIAHP by depolarizing pulses (200 ms @ 20 mV)^15^, which induce a large, slowly decaying outward tail current following the depolarizing pulse (Figure 1F), and changes in the area of sIAHP were evaluated. Astrocyte activation significantly increased the sIAHP area in CA1 pyramidal neurons of Gq-DREADD mice (Figure 1G), whereas no changes were found in control mice (astrocytes transfected with AAV5-GFAP-mCherry) (Figure 1G). To further confirm that astrocytic Ca^2+^ signaling activation impacts the slow component of IAHP in CA1 neurons, we selectively blocked the KCa3.1 channels responsible for sIAHP^49,50^. As expected, the sIAHP increase was prevented in the presence of TRAM-34 (5 µM) (Figure S1A).

The astrocyte-induced modulation of sIAHP was more evident when studying the kinetics of the conductance, as we observed significant changes in the exponential function (*τ*) of the slow but not the medium component (Figure S1B). Interestingly, the sIAHP potentiation was not abolished by the sodium channel antagonist tetrodotoxin (TTX; 1 µM), which prevents action-potential generation, indicating that this phenomenon did not depend on neuronal activity (Figure 1G) and suggesting a direct effect on the recorded neuron. To exclude indirect effects of CNO actions over the Gq-DREADD expressing astrocytes, we performed paired whole-cell recordings in neurons and astrocytes, and selectively ablated G-protein signaling by adding GDPβS (10 mM)^27^ into the astrocyte recording pipette. In this experimental condition, the activation of G protein signaling cascades was prevented in astrocytes and CNO perfusion failed to increase the sIAHP area (Figure 1H). Additionally, transgenic mice with type-2 IP3 receptors knocked out (IP_3_R2^-/-^ mice)^51^, in which IP3-mediated astrocytic Ca^2+^ signaling is largely abolished ^24,52–54^, were analyzed. Hippocampal astrocytes from IP_3_R2^-/-^ mice were transfected with Gq-DREADD as previously described, and, in contrast to control animals, no differences were found in the sIAHP area after CNO application (Figure 1H). Taken together, these results indicate that astrocytes from hippocampal CA1 region through Ca^2+^-dependent signaling modulate pyramidal neuron excitability by regulating sIAHP conductance.

### Astrocytes regulate sIAHP via A1 adenosine receptors

We hypothesized that the effect observed in neurons in response to astrocytic activation was directly induced by gliotransmitter release. Previous works indicate that metabotropic glutamate receptors play an important role in the regulation of the sIAHP and subsequent control of epileptiform activity^15^. Interestingly, the increase of sIAHP induced by selective astrocytic Gq-DREADD activation remained in the presence of metabotropic glutamate receptor antagonists MPEP (50 µM) and LY367385 (100 µM) in slices perfused with TTX (Figure 2B), ruling out a role of these receptors in sIAHP regulation by astrocytes. Accumulated evidence has demonstrated that astrocytes release ATP after intracellular Ca^2+^ increase, which could act upon P2-purinergic receptor or be then converted to adenosine by ectonucleotidases and activate A1Rs^46,47,55–58^. To determine if any of these receptors underlie sIAHP upregulation in pyramidal cells, suramin (10 µM) -a potent blocker of P2Y2 purinergic receptor-or CPT (1 µM) -a high affinity adenosine A1 receptor antagonist-were applied in the presence of TTX. We found that CPT but not suramin blocked the increase of sIAHP area (Figure 2A), indicating that astrocyte regulation of this conductance is mediated by A1Rs. As expected, both the decrease of firing frequency (Figure 2C) and the increase of spike-frequency adaptation (Figure 2D) induced by q-DREADD activated astrocytes were prevented when slices were perfused with the A1R antagonist CPT.

**Figure 2:**
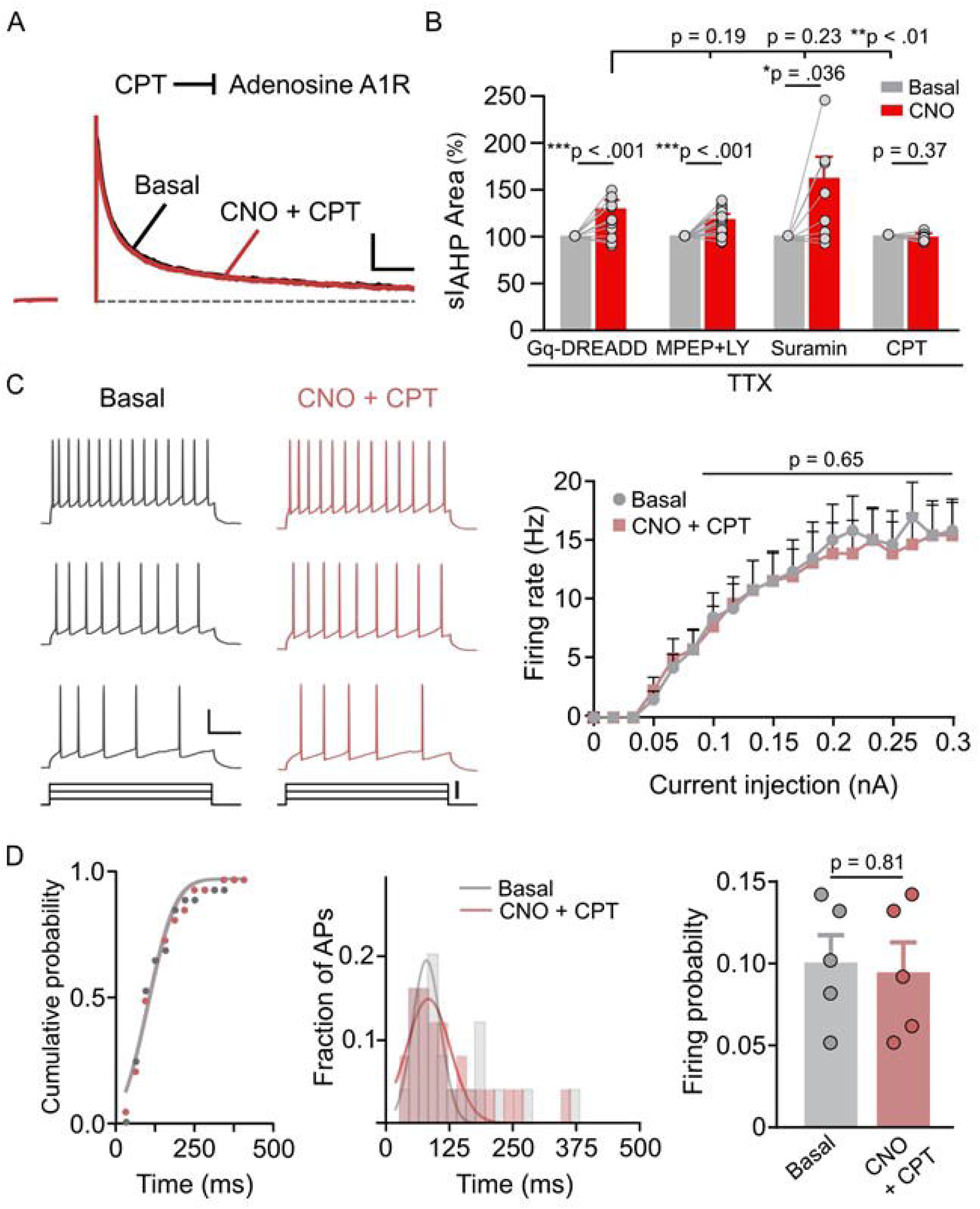
Neuronal activity is negatively controlled by adenosine modulation of sAHP. **A**, Representative traces of sIAHP recordings evoked before and after astrocyte stimulation with CNO 10 μM and blockade of adenosine A1 receptors by 1 μM CPT. Scale bar: 0.3 nA, 0.25 s. **B,** The increase in sIAHP by CNO perfusion -in presence of TTX-was not affected by adding mGluR5 (MPEP) + mGluR1 (LY367385) (n=23; Paired *t-*test) or P2Y2 (suramin; n=8; Paired *t-*test) receptor antagonists, but was markedly reduced by CPT (n=11; Mann-Whitney test). Significant differences with respect to Gq-DREADD were established by unpaired Student’s t-test or Mann-Whitney. **C,** Left: Traces of current-clamp recordings through increasing depolarizing current injection in Basal (black traces) and CNO + CPT (red traces) experimental conditions. Scale bar: 20 mV, 0.1 s. Right: Summary data show no decrease in pyramidal neuron firing frequency after astrocyte stimulation in presence of the adenosine A1R antagonist CPT. **D**, Astrocyte-driven neuronal spike accommodation changes seen after CNO perfusion are blocked by CPT (n=5; two-way ANOVA, F_(1, 160)_ = 0,2011; post hoc comparison with Bonferroni test) (KS test, P>0.5). Also, no changes were observed in the study of the probability of a neuron firing an action potential during the last 100 ms of the pulse. Data are represented as mean ± SEM.

To reinforce these findings, 1 mM adenosine was locally applied while both sIAHP (Figure 3A) and Ca^2+^ levels (Figure S2) were monitored in CA1 pyramidal neurons. In presence of TTX, local application of adenosine (1 bar, 2 s) induced an increase in the sIAHP area (Figure 3A) and neuronal Ca^2+^ (Figure S2) that were counteracted when CPT was perfused in the bath (Figure 3A; Figure S2B). A1Rs mediate a decrease in cAMP via G_i/o_^59^. G_i/o_ GPCR signaling has also been reported to activate PLC through the βγ subunit^60,61^. Therefore, we blocked G protein signaling in the recorded neurons by intracellular loading with GDPβS (10 mM)^27^ or PLC specific inhibitor U73122 (10 µM)^62–64^ in order to elucidate the contribution of these intracellular pathways for the sIAHP enhancement. In this experimental condition, adenosine-evoked sIAHP potentiation was prevented by both GDPβS and U73122 (Figure 3B), indicating that A1 GPCR signaling pathways increase Ca^2+^ by activation of PLC in agreement with previous results^65^. In line with these results, CNO perfusion in astrocytic Gq-DREADD mice failed to induce an increase in sIAHP when pyramidal neurons were intracellularly dialyzed with GDPβS or U73122 (Figure 3C). Interestingly, both the decrease in the frequency of action potential firing and the increase in the spike-frequency adaptation induced by CNO in astrocytic Gq-DREADD mice were blocked in the presence of GDPβS (Figure 3D) or U73122 (Figure 3E). Taken together, these results indicate that astrocytes, through ATP/adenosine release, induce a powerful tune of sIAHP current via GPCR-PLC intracellular pathway that regulates neuronal firing properties.

**Figure 3:**
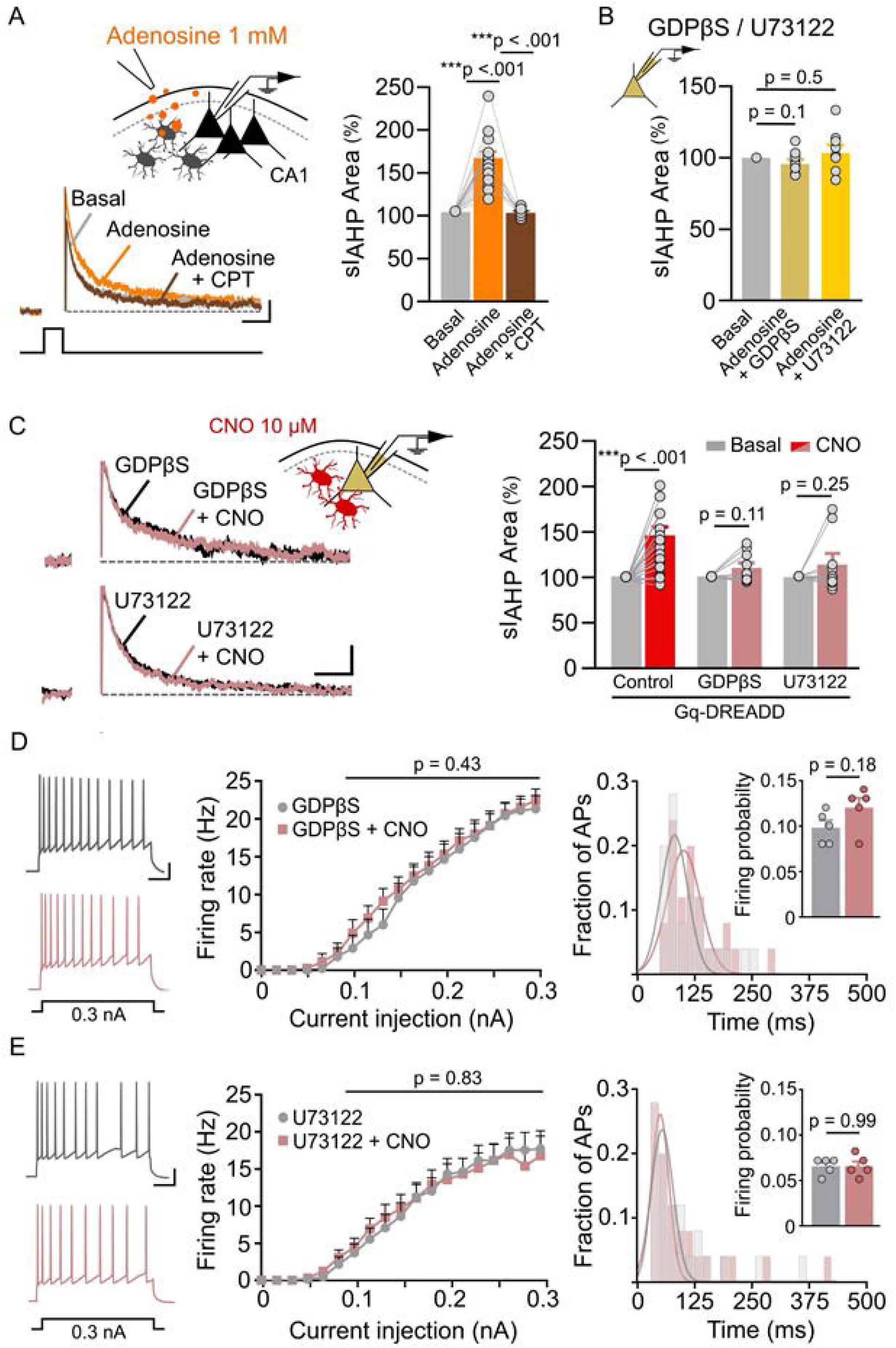
Astrocytes regulation of sIAHP and neuronal intrinsic excitability depends on G-proteins/PLC neuronal pathway signaling. **A,** Schematic for studying sAHP currents evoked by a local puff of adenosine 1mM in presence of TTX. Direct activation of A1Rs induced a rise in sIAHP (n=17) counteracted by CPT addition to the bath (n=8) (One-way ANOVA, F_(2, 39)_ = 51.42). **B,** No response was observed after adenosine application when blockade of G protein signaling (GDPβS; n=9) or inhibition of PLC cascade (U73122; n=10) was applied to the registered pyramidal neuron (Paired *t-*tests). **C**, As for **B**, but for astrocyte activation via CNO application (GDPβS; n= 11; U73122; n=11). **D**, **E**, Left: Representative traces of current-clamp (CC) recordings from Basal (gray traces) and CNO + GDPβS (red trace) or CNO + U73122 (red trace) experimental conditions in response to a depolarizing pulse (0.3 nA, 500 ms). Right: Intracellular application of GDPβS to block G-protein signaling or U73122 to specifically inhibit PLC, together with bath perfusion of CNO, prevented both spike firing frequency and accommodation modifications compared to the pretreatment control (n=9 for GDPβS; two-way ANOVA, F_(1, 320)_ = 0,61; post hoc comparison with Bonferroni test; KS test, P=0.47) (n=13 for U73122; two-way ANOVA, F_(1, 479)_ = 0,04; post hoc comparison with Bonferroni test; KS test, P>0.5). Likewise, the previously observed decrease in firing probability due to CNO perfusion for the last 100 ms of the pulse was no longer found in neurons filled with these drugs (n=5 for each condition; Paired *t-*tests). Data are represented as mean ± SEM.

### Astrocytic GABA_B_ receptors underlie sIAHP regulation

ATP/adenosine can be released by astrocytes under different scenarios, including pathological states^58,66–68^. Previous works indicate that GABAergic interneuron activity stimulates the release of ATP/adenosine by activation of astrocytic GABA_B_Rs^41,45,47^. Therefore, we hypothesized that astrocytic-dependent modulation of sIAHP in CA1 pyramidal neurons might be triggered by GABAergic interneuron activity. High-frequency stimulation (HFS) protocols of glutamatergic Schaffer-collateral pathway (SC) have been shown to induce interneuron GABA release and, consequently, GABA_B_R-mediated astrocytic Ca^2+^ and ATP/adenosine release that mediates heterosynaptic depression^41,69^. Therefore, we test this experimental approach to validate our hypothesis (Figure 4A). To isolate the effect mediated through GABA_B_ receptors exclusively, selective antagonists for GABA_A_, AMPA and NMDA antagonists (picrotoxin, D-AP5 and CNQX, respectively) were perfused to the hippocampal slices. To confirm that astrocytic Ca^2+^ signaling was recruited by HFS stimulation, cyto-GCaMP6f was injected in the hippocampal region to monitor intracellular Ca^2+^ levels in astrocytes, showing robust responses to HFS protocol^41^ (Figure 4B). As expected, HFS-induced astrocytic Ca^2+^ signaling was blocked by GABA_B_R antagonist CGP 55845 (CGP), but not by GABA_A_R antagonist picrotoxin (Figure 4C). In this condition, HFS evoked a decrease in the firing rate of CA1 pyramidal neurons (Figure 4D, E), which was prevented by adding CGP or CPT to the bath perfusion (Figure 4F). As anticipated, we observed that HFS elicited an increase of sIAHP area (Figure 4G, H). Such modulation was prevented when both GABA_B_Rs or A1Rs were pharmacologically blocked with CGP and CPT, respectively (Figure 4I). Remarkably, after ablation of G-protein signaling in CA1 neurons (Figure 4J), no sIAHP increment was detected after HFS.

**Figure 4.**
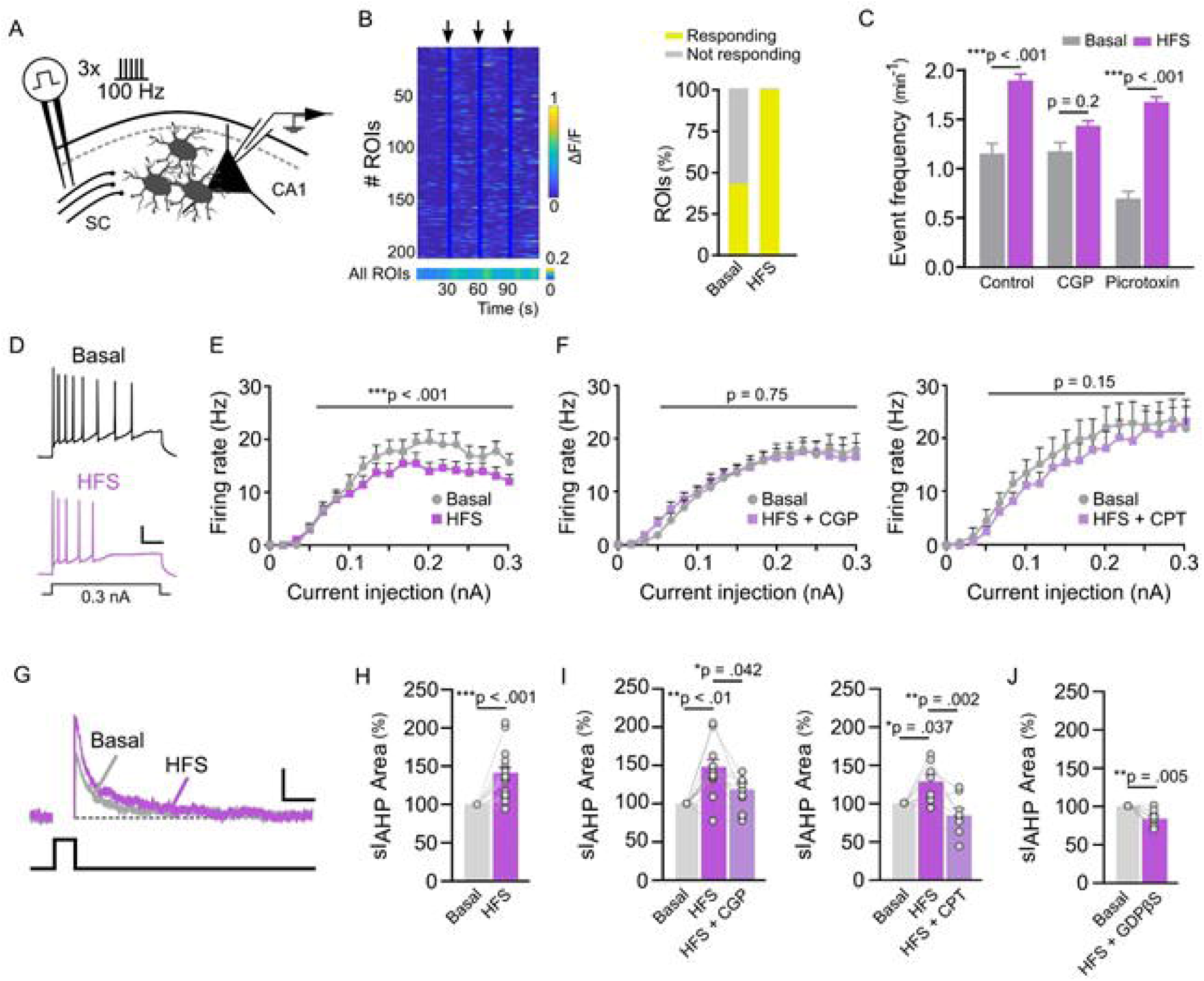
Intrinsic properties of pyramidal neurons can be modulated by electrical stimulation of hippocampal interneurons. **A**, Schematic drawing showing the electrical stimulation of Schaffer-collateral pathway (SC) through a high-frequency stimulation (HFS) paradigm (see Methods) and recordings from CA1 pyramidal neurons. **B**, Left: Raster plot of ROIs activity showing astrocyte Ca^2+^ levels in response to activation of interneurons - denoted by black arrows- (n=202). Right: Prior to interneuron stimulation with HFS, 86 astrocytes showed Ca^2+^ activity (Basal); their number increased to 201 after HFS application (HFS). **C**, The frequency of astrocytic Ca^2+^ events was increased after interneuron stimulation and was prevented by adding GABA_B_R antagonist CGP 55845 (CGP) to the bath perfusion (n=409; 18 slices from two mice; Mann-Whitney tests) but not in presence of GABA_A_R antagonist picrotoxin. **D,** Examples of firing patterns evoked after application of HFS protocol using 500 ms somatic current injection in current-clamp mode. Scale bar: 20 mV, 0.1 s. **E, F**, Plots of the firing responses versus stimulus intensity for the basal condition and after SC stimulation in absence or presence of GABA_B_R or A1R antagonists (n=19 for HFS; two-way ANOVA, F(1, 18) = 16.04) (n=10 for HFS + CGP; two-way ANOVA, F(1, 9) = 0,11) (n=6 for HFS + CPT; two-way ANOVA, F(1, 5) = 2,89). Post hoc comparisons performed with Bonferroni. **G**, Representative voltage-clamp recordings of the sIAHP in basal and after HFS stimulation of interneurons. Scale bar: 100 pA, 0.25 s. **H**, **I**, HFS of SC elicits an increase of sIAHP area (n=19; Paired t-test) that is disrupted if GABA_B_R in astrocytes or A1R in CA1 neurons are pharmacologically blocked with CGP and CPT, respectively (n=10 for HFS + CGP; one-way ANOVA, F(2, 28)=7.75) (n=8 for HFS + CPT; one-way ANOVA, F(2, 21)=8.6). **J**, Same stimulation protocol as in **H** but after ablation of G-protein signaling in pyramidal neurons (n=8; Paired t-test). Data are represented as mean ± SEM.

To further support the role of GABAergic interneuron-astrocyte cellular pathway in the control of CA1 pyramidal excitability, a more specific interneuron manipulation was used. Selective optogenetic stimulation was achieved by injection of ssAAV-1/2mDlx-HBB-hChR2-mCherry into hippocampal region^70,71^ (Figure 5A). Light stimulation of hippocampal interneurons caused an increase of sIAHP area in CA1 pyramidal neurons (Figure 5B, C) that was counteracted when GABA_B_R and A1R were pharmacologically blocked by CGP and CPT, respectively (Figure 5B, C). In addition, optostimulation of GABAergic interneurons in IP_3_R2^-/-^ mice, which show significant astrocytic Ca^2+^ impairments, did not evoke changes in sIAHP area (Figure 5D), confirming the astrocytic Ca^2+^-dependent mechanism that underlies this phenomenon. Furthermore, as found by endogenous activation of synaptic pathways (HFS experiments), optogenetic interneuron activation induced a decrease in the firing rate of CA1 pyramidal neurons (Figure 5E, F), which was blocked by adding CGP and CPT to the bath perfusion (Figure 5G). The selective astrocytic Ca^2+^ responses to optogenetic activation of GABAergic interneurons were confirmed by transfecting hippocampal region with both mdlx-ChR2 and gfaABC1D-GCaMP6f virus to target interneurons and astrocytes, respectively. Following optogenetic stimulation, astrocytes showed an increase in Ca^2+^ events (Figure 5H, I) that was prevented by GABA_B_R blockade (Figure 5J).

**Figure 5:**
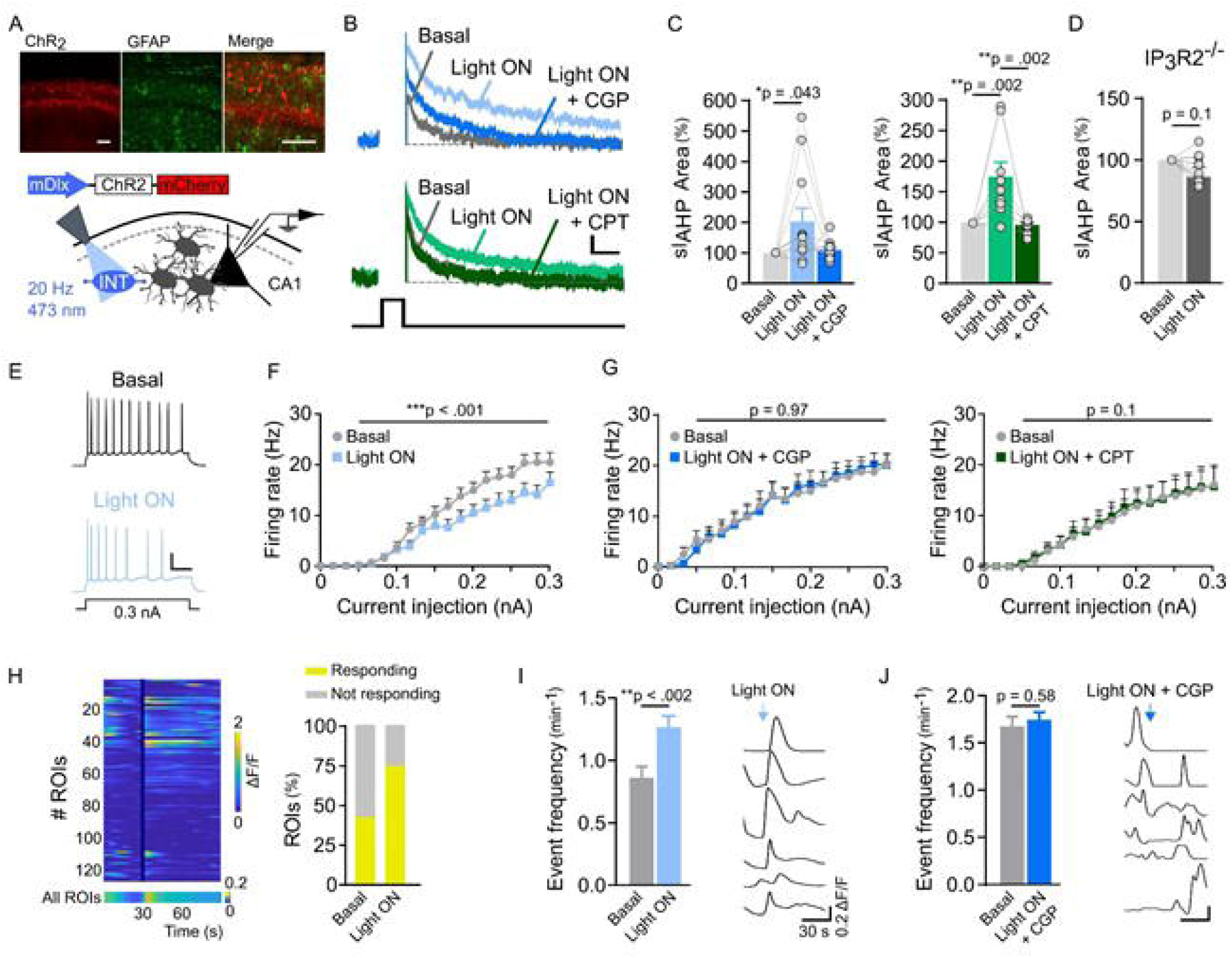
GABA released from interneurons limits neuronal excitability through astrocytic GABA_B_ and neuronal A1 receptors. **A**, Top: Confocal image of AAV1-1/2mDlx-HBB-hChR2-mCherry (red) expression in interneurons in the CA1 region of the hippocampus and immunocytochemical localization of astrocytes with GFAP staining (green). Scale bar: 50 μm. Bottom: Schematic drawing depicting the depolarization protocol used to optostimulate hippocampal interneurons (INT) (see Methods) to induce a similar heterosynaptic activation, in the presence of GABA_A_, AMPA and NMDA antagonists to isolate GABA_B_R astrocyte-mediated sIAHP potentiation. **B**, Representative voltage-clamp recordings of the sIAHP in basal condition, after interneuron stimulation (Light ON) and during superfusion with 1 μM CGP 55845 or 1 μM CPT. Scale bar: 100 pA, 0.25 s. **C**, Activation of hippocampal interneurons increases sIAHP in a GABA_B_R- and A1R-dependent manner (n=12 for CGP; n=9 for CPT; one-way ANOVA). **D**, Same stimulation protocol as in **A** but in IP_3_R2^-/-^ mice. The absence of sIAHP response provides evidence for the active involvement of astrocytes in the effect of interneurons upon the excitability of pyramidal neurons (n=10; Paired *t*-test). **E**, Examples of firing patterns evoked after light stimulation of interneurons using 500 ms somatic current injection in current-clamp mode. Scale bar: 20 mV, 0.1 s. **F**, **G**, Plots of the firing responses versus stimulus intensity for the basal condition and after interneuron optostimulation in absence or presence of GABA_B_R or A1R antagonists. Note the significant decrease in spike output of pyramidal neurons when no drug is applied (n=12 for Light ON; two-way ANOVA, F_(1, 220)_ = 120,5) (n=9 for Light ON + CGP; two-way ANOVA, F_(1, 160)_ = 0,00197) (n=10 for Light ON + CPT; two-way ANOVA, F_(1, 180)_ = 2,667). Post hoc comparisons performed with Bonferroni. **H**, To verify the role of astrocytes in this putative network we injected GCaMP6f to monitor astrocytic Ca^2+^ while triggering GABA release. Left: Raster plot of ROIs activity showing astrocyte Ca^2+^ levels in response to activation of interneurons (n=126). Right: Prior to interneuron stimulation with light, 53 astrocytes showed Ca^2+^ activity (Basal); their number increased to 94 after light application (Light ON). **I**, The frequency of astrocytic Ca^2+^ events was boosted after interneuron optostimulation -denoted by blue arrow- (n=126; 10 slices from three mice; paired *t-*test). **J**, Optogenetic activation of interneurons after blockade of GABA_B_ receptors failed to evoke changes on astrocytic activity (n=147; 11 slices from two mice; paired *t-*test). Data are represented as mean ± SEM.

Finally, to reinforce the impact of GABAergic interneuron-to-astrocyte signaling on sIAHP modulation and control of the CA1 neurons excitability, we generated transgenic mice combining *Gabbr1* (GABA_B1_)-floxed mice^71^ with viral expression of GFAP-Cre recombinase (AVV2.5-GFAP-Cre-mCherry) (Figure 6A) in CA1, leading to the specific downregulation of GABA_B_R in astrocytes (GB1-cKO mice; Figure S3). To assess whether neuronal intrinsic properties were altered by transgenic suppression of astrocytic GABA_B_ receptors, we examined cell excitability in pyramidal neurons of CA1 (Figure S4). Interestingly, we found that, with the same injected current, the firing frequency and the number of spikes in GB1-cKO pyramidal neurons were higher with respect to control ones. In addition, we injected both mDlx-ChR2 and GFAP-hM3Dq into hippocampal region to selectively activate GABAergic interneurons with light and astrocytes with CNO (Figure 6A). First, the analysis of sIAHP area on GB1-cKO slices indicated no effect on CA1 pyramidal neurons after interneuron optostimulation (Figure 6B, Light ON). Accordingly, ChR2-driven interneuron excitation failed to decrease the firing rate (Figure 6D). However, chemogenetic astrocytic stimulation in GB1-cKO evoked a robust enhancement of sIAHP area (Figure 6C) that promoted the decrease of excitability and increase of spike-frequency adaptation in CA1 pyramidal neurons (Figure 6E). These data confirm the role of astrocytes in tuning hippocampal CA1 excitability via GABA release from interneurons, revealing the key contribution of astrocytes in controlling hippocampal microcircuit activity.

**Figure 6:**
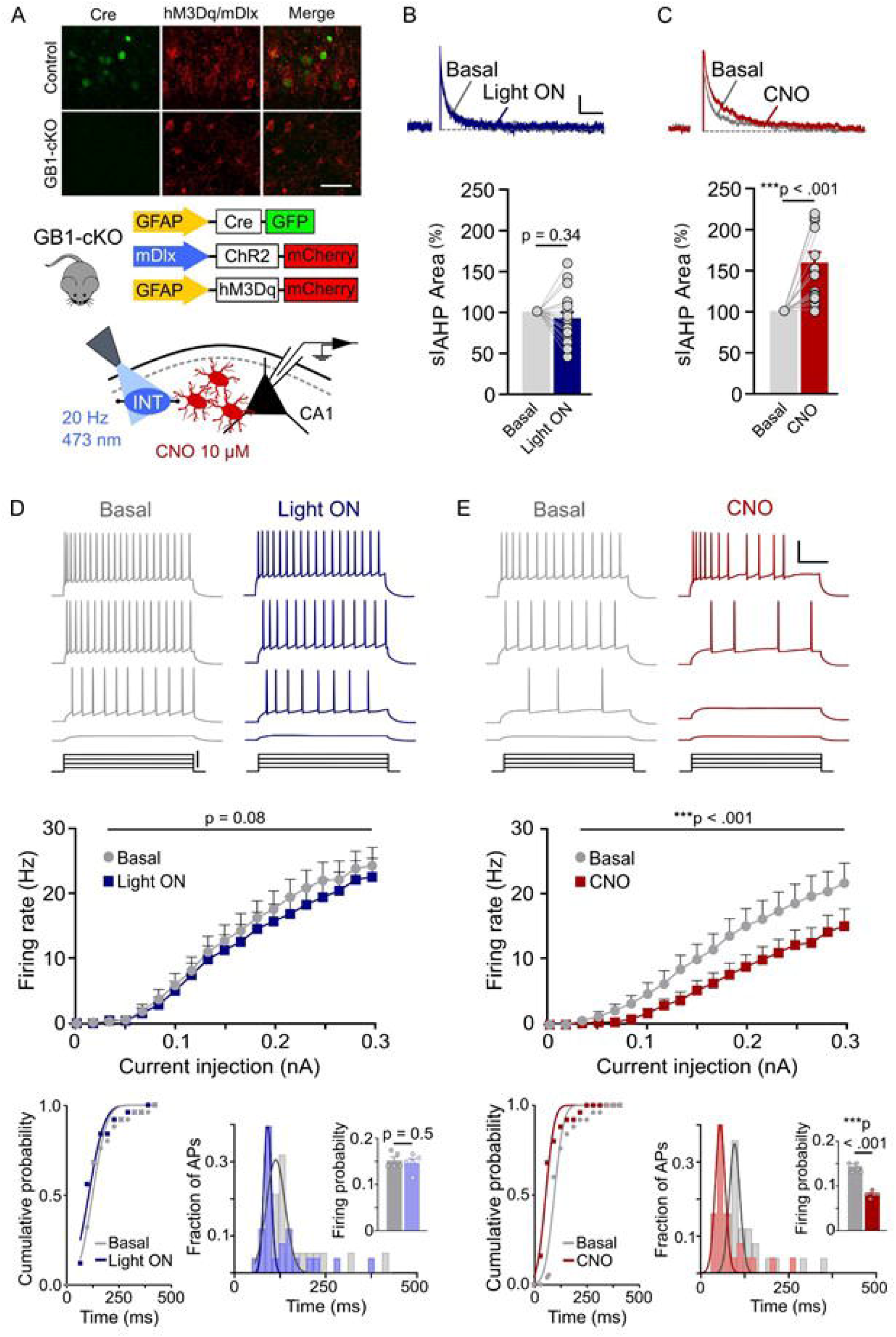
Interneuron modulation of intrinsic neuronal properties is disrupted by absence of astrocytic GABA_B_Rs. **A**, Top: Immunofluorescence image that confirms GABA_B_R downregulation in hippocampal astrocytes. Bottom: Scheme of sIAHP and firing rate recordings of pyramidal neurons surrounded by ChR2-positive interneurons (INT) and hM3Dq-positive astrocytes in an astrocytic GABA_B_R-negative background. The optogenetic protocol (30 s) was first applied to stimulate interneuron depolarization; afterwards CNO 10 μM was perfused in the bath to boost astrocyte activation. **B**, The sample voltage-clamp recording and mean area plot show no effect of the interneuron optostimulating protocol (Light ON) on the sIAHP-mediated current (sIAHP) from GB1-cKO mice (n=18; Paired *t*-test). Scale bar: 100 pA, 0.5 s. **C**, Direct CNO-treated astrocyte activation (CNO) in the GB1-null mice induces >50% potentiation in evoked sIAHP recordings from the same neurons in **B** (n=16; Paired *t*-test), revealing that astrocytic GABA_B_ receptors are necessary for sIAHP modulation by interneurons. **D**, Top: Opto- or chemogenetic stimulation of interneurons and astrocytes, respectively, from GB1-cKO mice elicits different firing frequency patterns through 500 ms somatic increasing depolarizing current injection. Note the expected linear increase in the number of action potentials in the Basal and Light ON conditions, in contrast to the decrease in firing frequency and increase in spike accommodation shown in the CNO condition (**E**), conferred by the larger sIAHP observed in **C**. Scale bar: 40 mV, 0.1 s, 0.3 nA. Bottom: Current-clamp recordings from pyramidal neurons of downregulated astrocytic GABA_B_R mice demonstrate the lack of effect of interneuron activation upon neuronal excitability. There are neither changes on spike output (n=14 for Light ON; two-way ANOVA, F_(1, 520)_ = 3.14; Post hoc comparisons with Bonferroni test) nor spike-frequency adaptation when light is applied (25 bins of 20 ms each; KS test, p= 0.28). The inset depicts that the firing probability for the last 100 ms of the pulse remains the same both in basal condition and after optogenetic manipulation (n=5; Paired *t-*test). **E**, On the contrary, neuronal excitability is modulated by direct astrocytic activation with CNO. Plot of the firing responses versus current injection for the basal condition and after CNO perfusion shows a significant decrease in the spike output (n=13 for CNO; two-way ANOVA, F_(1, 480)_ = 43.45; Post hoc comparisons with Bonferroni test) accompanied by an increase in spike-frequency adaptation (KS test, p < 0.001). Inset: The firing probability during the last 100 ms of the pulse declines significantly as only a small number of currents are initiated because of spike-accommodation (n=5; Paired *t-*test). Data are represented as mean ± SEM.

## DISCUSSION

The pivotal role of astrocytes in brain homeostasis, providing trophic, structural and metabolic support for neurons has been reported in the last decades^72–74^. Astrocytes are known to express receptors for different neurotransmitters, such as glutamate, GABA, endocannabinoids, dopamine, serotonin, ATP/adenosine, acetylcholine or opioids^21–24^ and have the ability to sense multiple neuronal signals conferring a high variety of astrocytic responses^75–77^. In hippocampal region, astrocytes are well known to release glutamate, ATP and GABA^28,30,46,47,78,79^. In this scenario, we hypothesized that some of these gliotransmitters could regulate the sIAHP and, consequently, modify neuronal excitability. sIAHP is a Ca^2+^-induced potassium current that underlies action potential firing pattern modulation^80,81^, and provides a strong negative feedback to control neuronal excitability in physiological^12^ and pathological conditions^13–15^. The present study uncovers how astrocytes modulate membrane intrinsic properties of CA1 pyramidal neurons through regulation of sIAHP. Hence, by modulating this current, astrocytes are indeed in a critical position to regulate neuronal excitability, but also hippocampal network activity.

Using pharmacological and chemo-stimulation approaches, we indeed found that ATP/adenosine released by astrocytes increases sIAHP conductance and, consequently, decreases firing rate via adenosine A1Rs. We first demonstrated that Gq-DREADD astrocyte activation enhanced the area of the sIAHP, even when TTX is bath perfused, ruling out a neuronal action potential-dependent activity. Remarkably, sIAHP recovers to baseline after addition of specific adenosine A1R antagonist CPT, but not in the presence of suramin, a purinergic receptor antagonist. However, it is possible that ATP release can stimulate the same response since it is promptly converted to adenosine by extracellular ectonucleotidases^58,82–84^. Although with relevant roles in synaptic modulation^29^, glutamate released by astrocytes was not primarily involved in the regulation of sIAHP, as we found that modulation of sIAHP was not affected in presence of specific metabotropic glutamate receptor antagonists^15^. Previous studies have shown that adenosine upregulates sIAHP, although the specific mechanisms, both intracellular and intercellular, underlying this effect remained poorly understood^85,86^. Here, we show that activation of A1 receptors, which couple to heterotrimeric G-proteins of the G_i/o_ family, increases sIAHP in CA1 pyramidal cells through a PLC-induced Ca^2+^ mobilization-dependent mechanism. In addition, this effect was abolished by application of U73122 that inhibits inositol 1,4,5-trisphosphate (IP_3_)^25,28,87^ increment and Ca^2+^ release from the endoplasmic reticulum^88^. Previous evidence points to KCa3.1 as the Ca^2+^-dependent K^+^-channel underlying control over membrane excitability through most of the sAHP component not only in hippocampus^50,89^, but also in the neocortex^90^. In this line, we found that astrocyte activation was unsuccessful to increase the sIAHP in the presence of TRAM-34, a selective blocker of KCa3.1 channels^49,50^ confirming the roll of this conductance in the observed phenomena.

Astrocytes can be activated by hippocampal interneurons triggering multiple cellular responses^30,91,92^. GABA_B_ receptor activation has been shown to trigger intracellular Ca^2+^ elevations in astrocytes^93^, which is sufficient to evoke astrocytic ATP release, being degraded to adenosine^83^, when is triggered by electrical HFS of glutamatergic SC^41,45–47^. Therefore, our study assessed the critical role of interneuron-astrocyte signaling pathway, through GABA_B_Rs and A1, to the regulation of sIAHP and firing rate of CA1 pyramidal neurons. We found that both electrical HFS of SC and optostimulation of interneurons were very effective eliciting positive responses in the astrocytic Ca^2+^ signals and modulating intrinsic excitability by enhancing sIAHP. However, no effect was observed in optogenetic experiments on adult mice IP_3_R2^-/-^ or animals with GABA_B_R-specific astrocytic deletion^71^. Interestingly, direct astrocyte activation by chemogenetic stimulation effectively leads to a decrease in CA1 cell excitability and sIAHP increase, which provides evidence for a functional unit for the control of neuronal activity that relies on interneuron/astrocyte/pyramidal neuron interactions. Taken together, we propose a coordinated heterosynaptic circuit that involves GABAergic interneuron activity, astrocytic GABA_B_R activation and ATP/adenosine release, and activation of A1 in CA1 pyramidal neurons that downregulates the intrinsic excitability by boosting sIAHP. Our findings reveal a new scenario where interneurons play a dual control over CA1 since directly inhibits synaptic inputs by GABA_A_ receptors and, simultaneously, modifies intrinsic excitability by astrocytic GABA_B_-mediated ATP/adenosine release. Thus, although CA1 cells receive great glutamatergic excitation, their excitability is critically controlled by interneurons, increasing the stability of hippocampal network and allowing proper coding^94^. Therefore, the fact that astrocytes are involved in regulation of CA1 excitability extends our knowledge about the functional operation of hippocampal circuits in health and disease. In this scenario, astrocytes emerge as key elements to understand hippocampal-related disorders associated with disrupted neuronal activity.

## Data and code availability

- This paper does not report original code.
- Any additional information required to reanalyze the data reported in this work paper is available from the lead contact upon reasonable request.

## Acknowledgements

We gratefully acknowledge Dra. Gertrudis Perea for generously donating *Gabbr1*-floxed and IP3R2^-/-^ mice, the permanent work feedback and critical reading of the manuscript. This work was supported by Grant PID2020-116327GB-I00, MCIN/AEI/10.13039/501100011033, Spain to E.D.M.

## Author contributions

Conceptualization E.D.M.; Methodology, E.D.M; Investigation, S.E. and S.A.; Formal Analyses, S.E. and S.A; Writing – Original Draft, S.E. and S.A.; Writing – Review and Editing, S.E., S.A and E.D.M; Visualization, S.E., S.A and E.D.M; Project administration, E.D.M.; Funding Acquisition E.D.M.; Resources, E.D.M.; Supervision, E.D.M.

## Methods

### Animals

Mice were housed under 12/12-h light/dark cycle and up to four animals per cage. This study was carried out in 2-3 month old female and male C57BL/6J, *Gabbr1*-floxed and IP3R2^-/-^ mice (generously donated by Dr. G. Perea). All procedures were performed in accordance with the guidelines from European Union Council Directive (86/609/European Economic Community).

### Stereotactic Surgery

Two months old mice were anesthetized with isoflurane (1.5–2% in O_2_) and placed in a stereotactic frame (Kopf Instruments). The skull was exposed, and a small craniotomy was performed. Viral vectors were injected bilaterally using a Hamilton syringe attached to a 29-gauge needle at a rate of 0.05 µl/min. All viral constructs were delivered into the CA1 region of the hippocampus: anterior-posterior (AP): -2.65 mm; medial-lateral (ML): +/- 2.2mm; dorsal-ventral (DV): -1.95 mm. After infusion, the needle was kept at the injection site for 10 minutes and then slowly withdrawn before the incision was sutured. To monitor astrocytic Ca^2+^ signaling we used the genetically encoded Ca^2+^ indicator (GECI) AAV5-gfaABC1D-cyto-GCaMP6f (Addgene 52925) (viral injections of 0.6 µl) (viral titer 1.3×10^13^ viral genomes (vg) per ml). Transgenic mice to determine the impact of GABAergic astrocyte signaling on hippocampus were generated by combining adult *Gabbr1* (GABA_B_)-floxed mice with expression of 0.5 µl GFAP-Cre recombinase (AAV2.5-GFAP-Cre-mCherry) (UNC Vector Core) (viral titer 4.3×10^12^ vg per ml) in the hippocampus (GB1-cKO mice). We use Designer Receptors Exclusively Activated by Designer Drugs (DREAADs) approach to selectively activate astrocytes in CA1 hippocampal area via Clozapine-N-Oxide (CNO)^95^. To target astrocytes with chemogenetics we injected 0.60 µl bilaterally of the AAV5-GFAP-hM3D(Gq)-mCherry (Addgene 50478) (viral titer 2×10^13^ vg per ml) and the AAV5-GFAP-mCherry (for control experiments) (UNC vector core) in wild-type and GB1-cKO mice. GABAergic interneuron targeting was achieved with 0.45 µl of AAV1-1/2mDlx-HBB-hChR2-mcherry (Addgene 83898) (viral titer 7×10^12^ vg per ml) in wild-type and GB1-cKO mice. Mice were used 2 weeks after stereotactic surgeries.

### Mouse genotyping

We checked the presence of the knockout allele following Cre-mediated deletion of exon 7 and exon 8 in WT and GABA flox^+/+^ mouse cortical slices. DNA bands were detected after genomic DNA was extracted using an Extracta DNA prep for PCR—tissue kit (Quanta Bio) with primer number 322 (5′-CCATTGGACTCAGCTAATCCC-3′), 323 (5′-CTCCAGAGGTCTTGTTCAAGG-3′), 324 (5′-TGATCGGAATTCCTCGACTGC-3′) and 35 cycles of PCR (94°C for 15 s, 60°C for 15 s and an extension of 72°C for 1 min) with an Accustart II Gel Track PCR Supermix kit (Quanta Bio) was used. DNA fragments of 350 bp were generated for the wild-type allele (primer numbers 323 and 324) and 385 bp for the floxed allele (primer numbers 322 and 323) (Figure S3).

### Hippocampal slice preparation

Coronal hippocampal slices (350 µm) were made from 10-11 weeks old mice. Animals were decapitated and their brains rapidly sliced in ice-cold artificial cerebrospinal fluid (ACSF) using a vibratome (Vibratome Series 3000 Plus, St. Louis). Slices containing the hippocampal formation were incubated for 1 h at room temperature (21–24 °C) in ACSF that contained 124 mM NaCl, 2.69 mM KCl, 1.25 mM KH_2_PO_4_, 2 mM MgSO_4_, 26 mM NaHCO_3_, 2 mM CaCl_2_ and 10 mM glucose, and was gassed with 95% O_2_, 5% CO_2_, pH 7.3. Slices were then transferred to an immersion recording chamber and superfused at 2 ml/min.

### Electrophysiology

Electrophysiology recordings of CA1 pyramidal neurons and *Stratum Radiatum (SR)* interneurons were made in whole-cell configuration of the patch clamp technique. Patch electrodes were fabricated from borosilicate glass capillaries with a Brown-Flamming model P-80 micropipette puller (Sutter Instruments, Novato, CA) with resistances of 4–10 MΩ when back-filled with an internal solution. For recordings of the Ca^2+^-dependant K^+^-current (IAHP), the internal solution used contained (in mM): 150 KMeSO_4_, 10 HEPES, 4 ATP-Na_2_ (added immediately before filling) and 0.4 GTP-Na_2_, and was buffered to pH 7.2–7.3 with KOH. For spike frequency recordings, the internal solution contained (in mM): 135 KMeSO_4_, 10 KCl, 10 HEPES, 5 NaCl, 5 EGTA, 10 phosphocreatine, 2.5 ATP-Mg^+2^, and 0.3 GTP-Na^+^ (pH = 7.3). In some experiments, GDPβS (10 mM) or U73122 (10 µM) were included in the patch pipette to prevent G protein-mediated intracellular signaling in neurons or astrocytes or to study specific neuronal inhibition of phospholipase C, respectively. Also, TRAM-34 (5 µM) was added to the pippete solution when studying selective blockade of KCa3.1 channels. Electrophysiological recordings were obtained from one neuron per slice and 4-5 slices per mouse using a Multiclamp 700B amplifier and analyzed using a digital system (pClamp version 11.2, Molecular Devices). Series and input resistances were monitored throughout the experiment using a -5 mV pulse. Experiments were performed at controlled temperature (32-36°C). Cells were voltage-clamped at −65DmV and discarded when series and input resistances changed >20%. Signals were fed to a Pentium-based PC through a DigiData 1550B interface board and filtered at 1 KHz while acquired at 10 KHz sampling rate.

In all voltage clamp (VC) IAHP experiments, CA1 pyramidal cells were depolarized to -20 mV (to favor the increase of Ca^2+^ inside the cell) to evoke IAHP by voltage command pulses (200 ms) to 20 mV every 60 seconds^15^. The IAHP magnitude was quantified from the area under the current trace^15^. The baseline period includes three IAHP recordings. To prompt general astrocyte activation, CNO (10 µM) was perfused in the ACSF. In all the experiments GABA_A_, AMPA and NMDA receptors were inhibited by picrotoxin (50 µM), CNQX (20 µM) and D-APV (50 µM), respectively. Other compounds dissolved in ACSF were MPEP hydrochloride (50 µM), LY-367385 (100 µM), CPT (1 µM), suramin (10 µM), CGP 55845 hydrochloride (1 µM) and tetrodotoxin (TTX, 1 µM). Excitability was studied in current clamp (CC) mode using 500Dms somatic current injection in the absence of TTX.

### GABAergic interneuron stimulation

GABAergic interneurons were stimulated by two different methods. The high frequency stimulation (HFS) paradigm consisted in the positioning under visual guidance of a bipolar tungsten wire electrode connected to a stimulator and isolation unit (A-M Systems, Isolated Plus Stimulator, Model 2100). The stimulating electrode was placed in the *SR* near the CA1 pyramidal neurons to stimulate Schaffer collaterals. This tetanic stimulation consisted in the application of three trains at 100 Hz for 1 s delivered every 30 s with a stimulation intensity of 7 volts (V). We evoked IAHP 100 ms after each train and averaged three area traces.

The optostimulation of GABAergic interneurons was achieved by delivering blue light pulses at 473 nm and 20 Hz for 30 seconds (10 ms light pulses) directly over the CA1 hippocampal region. For that purpose, we used a light-emitting diode (LED; Doric Lenses, Québec, QC, Canada) and the Doric Neuroscience Studio Software to generate the illumination sequence required. In both cases, TTX and CGP 55845 hydrochloride were not added to the perfusion unless indicated.

### Calcium imaging

Calcium levels in proximal processes and soma of the astrocytes located in the *SR* of the CA1 region of the hippocampus were monitored by the calcium indicator AAV5-gfaABC1D-cyto-GCaMP6f. In the set of chemogenetic experiments, GCaMP6f was coinjected with AAV5-GFAP-hM3D(Gq)-mCherry to verify astrocytes responded to CNO. Astrocytes were imaged using epifluorescence microscopy with a CCD camera (ORCA-235; Hamamatsu, Japan) attached to the microscope (Olympus BX51WI) and illuminated during 200 ms with a xenon lamp at 490 nm. Images were acquired every 2 s. Areas with astrocytes showing both mCherry and GCaMP6f positive labeling were selected for analysis. Then, baseline was recorded for 30 s, followed by the application of a local puff of CNO (1 mM) -to transiently activate astrocytic DREADD receptors-for 60 s. CNO was delivered by pressure pulses through a micropipette (2 s air puff, 1 bar; Picospritzer II, Parker Hannifin, Mayfield Heights, OH, USA) in the presence of TTX. Regions of interest (ROIs) whose Ca^2+^ signal changed >2 standard deviations from baseline were selected from the recorded image. Ca^2+^ variations were quantified as changes in the fluorescence signal (F) over the baseline ΔF/F_0_ = ((F_t_-F_0_)/F_0_) and recorded from the astrocyte soma and proximal processes. Local Matlab software (MATLAB R2016; Mathworks Natick) custom-written plugin was used for computation of fluorescence values of each region of interest (ROI). Additionally, experiments with GCaMP6f coinjected with control virus (expressing the reporter gene, mCherry) were performed in order to distinguish between the responses due to spontaneous activity from those mediated by DREADDs. To reduce the effects of GABA_A_-mediated inhibitory synaptic transmission, Ca^2+^ experiments were performed in the presence of 50 µM picrotoxin.

### Immunohistochemistry

Mice were anesthetized and perfused through the left cardiac ventricle with 4% paraformaldehyde (PFA) in 0.1 M phosphate buffered saline (PBS) (pH 7.4). Brains were removed, post-fixed overnight (o/n) and then cut into 30 μm coronal sections using a vibratome. The slices were blocked for 60 min with PBS containing 5% normal goat serum (NGS) and 0.2% Triton X-100 at room temperature (RT), and incubated o/n at 4°C with the primary antibodies (Rabbit-antiGFAP: MAB3402, Merck Millipore, 1:500; Mouse-antiNeuN: MAB377, Merck Millipore, 1:500) diluted in the blocking solution. On the next day, slices were washed three times for ten minutes each in 0.2% Triton X-100 and PBS 0.1 M and incubated with the secondary antibodies for 2h at RT in the secondary antibody buffer (0,2% Triton X-100, PBS 0.1 M, 1% NGS). The secondary antibodies that were used for detection were the following: Alexa Fluor™ 488 donkey-antirrabit (*Invitrogen*, 1:500) against glial fibrillary acidic protein (GFAP) as a marker for astrocytes; and 405 goat-antimouse (*Invitrogen*, 1:500) against neuron-specific nuclear protein (NeuN) as a marker for neurons. Finally, sections were washed 3 times with PBS for 10 min and mounted in a fluorescent mounting medium (DABCO, Fluka). The stained sections were examined by laser confocal microscope (Leica).

### Drugs and chemicals

6-cyano-7-nitroquinoxaline-2, 3-dione (CNQX), D(2)-2-amino-5-phosphonopentanoic acid (D-AP5), (2S)-3-[(1S)-1-(3,4-Dichloro-phenyl)ethyl] amino-2-hydroxypropyl] (phenylmethyl) phosphonicacidhydrochloride (CGP 55845), 2-Methyl-6-(phenylethynyl) pyridine hydrochloride (MPEP hydrochloride), Octahydro-12-(hydroxymethyl)-2-imino-5,9:7,10a-dimethano-10aH-[1,3]dioxocino[6,5-d]pyrimidine-4,7,10,11,12-pentol (tetrodotoxin), 8,8’- [Carbonylbis[imino-3,1-phenylenecarbonylimino (4-methyl-3,1-phenylene) carbonylimino] bis-1,3,5-naphthalenetrisulfonic acidhexasodium salt (suramin hexasodium salt), 1-[6- [[(17β)-3-Methoxyestra-1,3,5(10)-trien-17-yl]amino]hexyl]-1H-pyrrole-2,5-dione (U73122) and 8-Chloro-11-(4-methyl-4-oxido-1-piperazinyl)-5*H*-dibenzo[*b*,*e*][1,4]diazepine (CNO) were purchased from Tocris. 8-Cyclopentyl-1,3-dimethylxanthine (CPT), Picrotoxin, Guanosine 5′-[β-thio]diphosphate trilithium salt (GDPßS), 9-β-D-Ribofuranosyladenine (adenosine) and (S)-(+)-a-amino-4-carboxy-2-methylbenzeneacetic acid (LY-367385) were purchased from Merck.

### Quantification and statistical analysis

The data were displayed, acquired and analyzed with the pClamp 11 software (Molecular Devices). Statistical analyses were performed using IBM SPSS Statistics version 27.0 and MATLAB (MATLAB R2016; Mathworks Natick). Data are expressed as the meanD±Dstandard error of the mean (SEM). We performed normality tests before applying statistical comparisons and assumptions of homogeneity were also checked. They were made using parametric Student’s t tests, one or two-way ANOVA and nonparametric tests when appropriate, and were followed by *post-hoc* comparisons with Bonferroni or Tukey as deemed appropriate. Unless indicated, two-tailed, unpaired or paired tests were used. p value and test employed are reported in the text and/or figures legends. Statistical differences were established with pD<D0.05 (*), pD<D0.01 (**), and pD<D0.001 (***). Sample size was based on values previously found sufficient to detect significant changes in hippocampal synaptic parameters in the past studies from the lab and randomization was not employed.

