## Supplemental information for "Astrocytes tune neuronal excitability through the Ca^2+^-activated K^+^ current sIAHP"

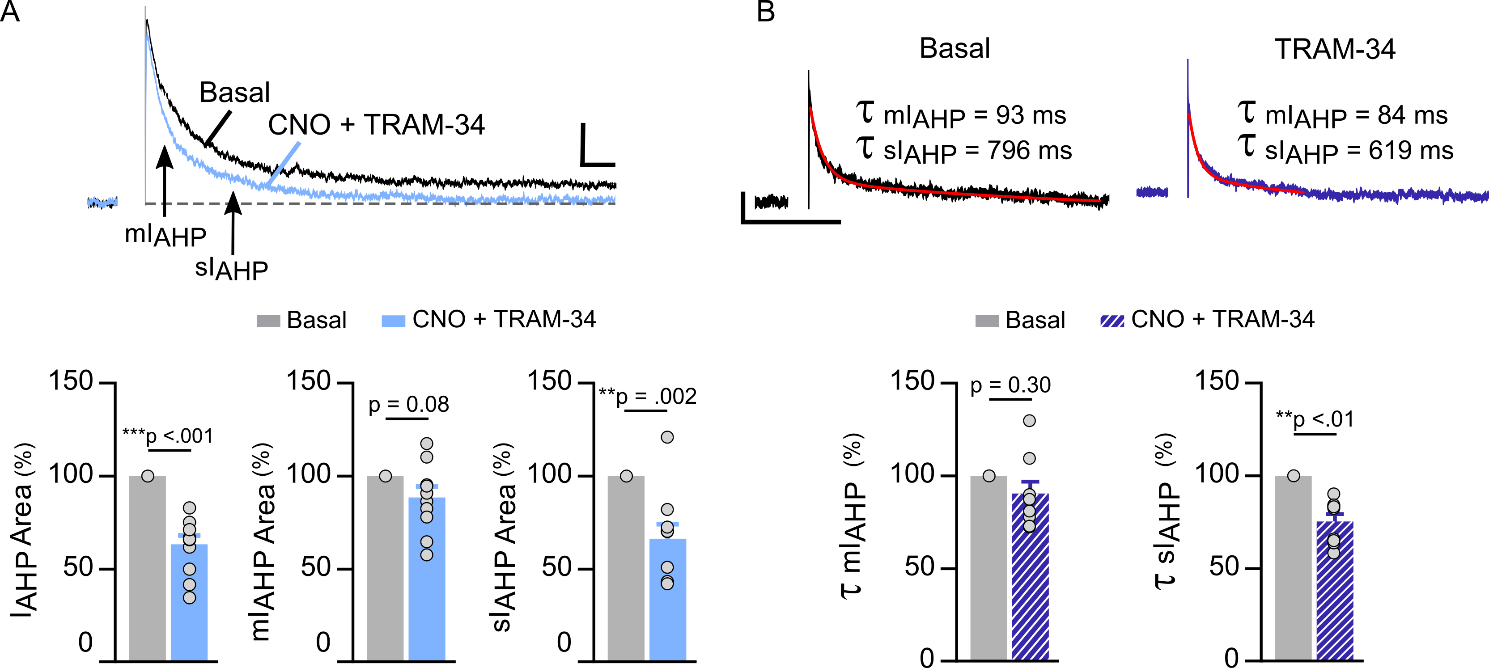

**Figure S1. KCa3.1 channels underlie control over membrane excitability. A**, Example of sAHP currents evoked by astrocyte activation before and after application of CNO with 5 μm TRAM-34. Scale bar: 100 pA, 0.25 s. KCa3.1 currents blockade decreases total and slow IAHP area -measured at 500 ms post-stimulus- (n=10; Paired *t-*tests). **B**, As the recordings show a decaying outward tail current (Top traces), we fitted a two term exponential function to the IAHP to study CNO-associated changes upon the medium and slow components of the IAHP (n=9; Mann-Whitney test). Data are represented as mean ± SEM.

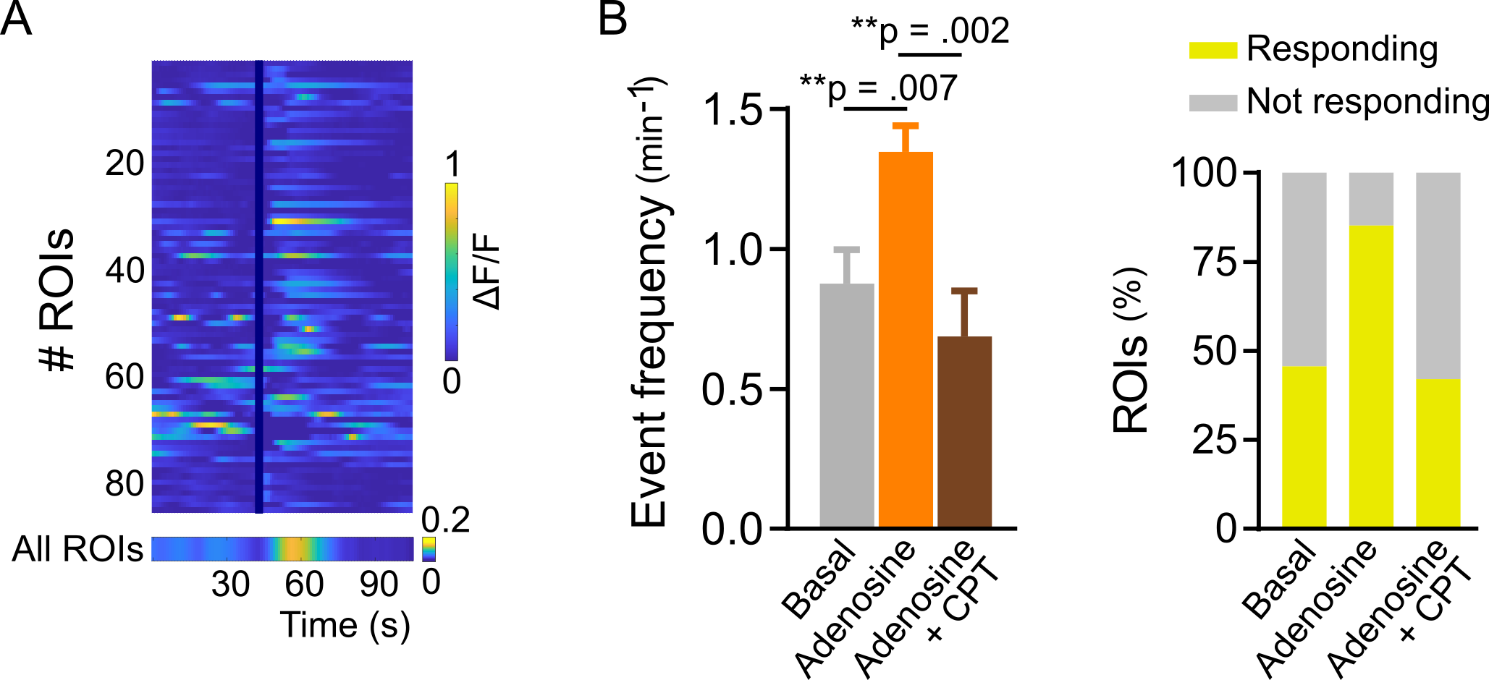

**Figure S2. Adenosine triggers A1-dependent Ca^2+^ signals in neurons. A**, Representative raster plot of ROIs activity showing neuronal Ca^2+^ levels, color coded according to fluorescence change (n=81). **B**, Left: Analysis of neuronal Ca^2+^ fluctuation properties showing mean responses for event frequency (9 slices from two mice) (One-Way ANOVA F_(2, 197)_ = 7.62; post hoc comparison with Bonferroni test). Right: Percentage of ROIs showing any event during the 30 seconds prior to application of adenosine (Responding ROI) (n=42), after application of 1 mM adenosine (n=69) and after addition of CPT to the bath (n=37). Data are represented as mean ± SEM.

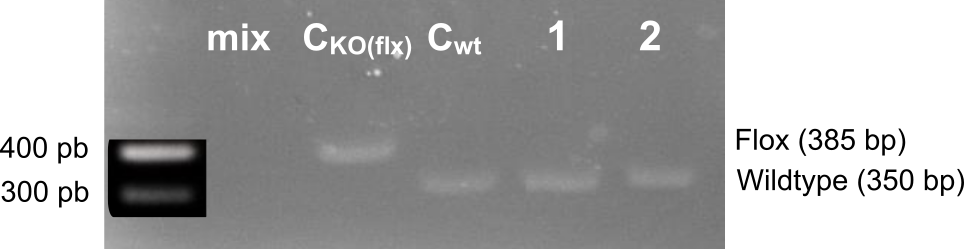

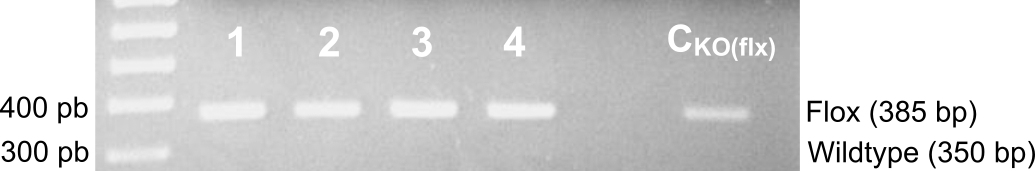

**Figure S3: Selective downregulation of GABA_B1_ in hippocampal astrocytes.** Quantitative PCRs of genomic DNA isolated from hippocampal region of slices from control (GABA_B_ flox^-/-^) and GABA_B_ flox^+/+^ mice show that GABA_B_ flox^+/+^ are positive for the targeted DNA sequence (allele of 385 bp; Flox) (n=6 slices from 2 mice for Control condition; n=17 slices from 4 mice for GABA_B_ flox^+/+^ condition).

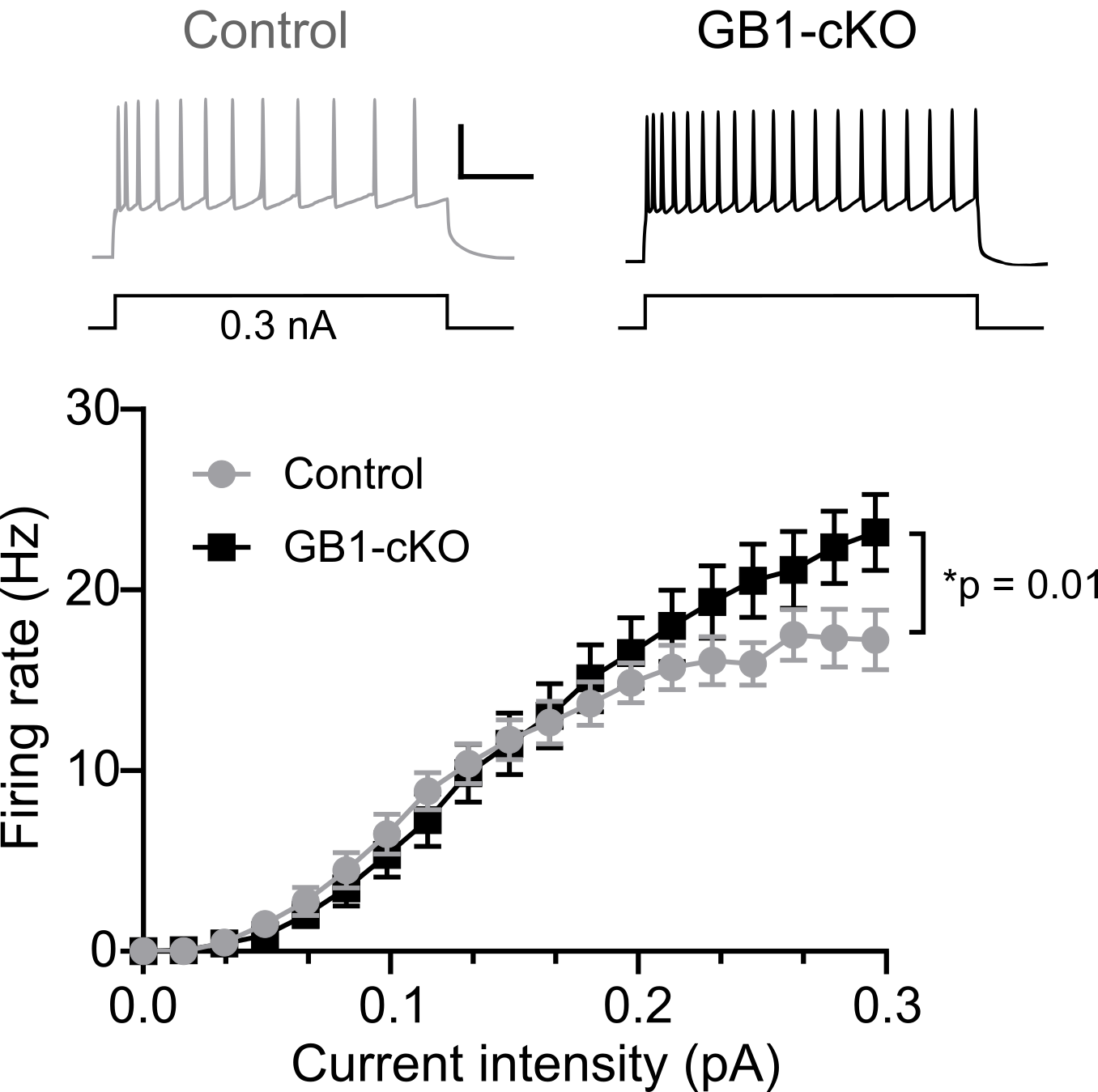

**Figure S4: Increased neuronal excitability of pyramidal neurons in the GB1-cKO mice.** Top: Examples of responses of neurons from control or transgenic mice to 0.3 nA of current injected for 500 ms in CA1. Scale bar: 20 mV, 0.1 s. Bottom: Graphs depict that the firing responses to different amounts of injected current are higher in GB1-cKO animals (n=20 for both conditions; two-way ANOVA, F(1, 920) = 6.41). Data are represented as mean ± SEM.
